# Assessing soluble and insoluble calcium sources for growth, biofilm formation, and biomineralization in *Bacillus subtilis*

**DOI:** 10.64898/2026.05.12.724540

**Authors:** Daniel Tchelet, Avichay Nahami, Adi Ioshpe Prem, Anand Murugan, Igor Lapsker, Yuval Dorfan, Ilana Kolodkin-Gal

## Abstract

Biofilms formed by soil microbes hold immense potential for bioremediation, carbon dioxide sequestration, and the development of sustainable cementitious materials. However, quantifying the complex temporal coupling among bacterial growth, extracellular matrix (ECM) production, and mineralization dynamics remains a significant challenge due to the inherent nonlinearity of these processes and signal noise in high-throughput assays.

To address this, we utilized an automated kinetic framework to evaluate the biomineralization competence of *Bacillus subtilis* under varying calcium regimes. Our results demonstrate that calcium carbonate promotes microbial growth as effectively as soluble calcium acetate, indicating that *B. subtilis* actively solubilizes the crystalline powder. Despite this growth efficacy, calcium carbonate was an inadequate source for macro-calcite production compared to organic salts. By quantifying expression from the sinI promoter, which initiates SinI production to activate matrix synthesis, we suggested that calcium-acetate-driven extracellular matrix (ECM) expression significantly enhances the structural template required for robust biomineralization. Kinetic expression analysis and consolidation assays indicate that overproduced ECM partially mitigates crystalline calcite growth defects, offering a baseline for utilizing mineral-rich construction waste

**Importance:** Biomineralization in B. subtilis has been studied as a potential avenue for developing engineered living materials. However, organic calcium salts, such as calcium acetate and calcium lactate, typically yield higher bioconversion efficiencies than certain inorganic sources often used in construction. This advantage has been largely attributed to the metabolic utilization of the organic anion as a carbon source, thereby promoting local alkalinity. Here, our experiments indicate that this limitation is influenced by structural and regulatory constraints rather than purely chemical barriers. Moreover, we explored a targeted synthetic biology approach to address this by genetically decoupling matrix production from native environmental sensing, thereby engineering strains with increased matrix production that can partially overcome the limitations of non-inductive substrates. This approach can contribute to the development of sustainable construction technologies, moving toward actively programmed Engineered Living Materials (ELMs).

## Introduction

*B. subtilis* has been termed the Swiss army knife in science and biotechnology (1) because of its unique position in the scientific field. *B. subtilis* is among the best understood organisms and the longest studied bacterium since its discovery in 1835 by Christian Gottfried Ehrenberg. It is not pathogenic but can form robust, heat-resistant spores, as well as form robust 3D communities, also designated biofilms (2).

Bacterial biofilms present a dominant multicellular mode of life for unicellular organisms (3). Their significance is difficult to underestimate, as they provide bacteria with increased resistance to environmental stressors and antimicrobials (4–8), enhance nutrient utilization through metabolic cooperation (9), support the transition into a dormant state and persistence (10, 11), intensify genetic information exchange and microbial diversity (12–14)Generate the environment where multiple species form a community (15) and, overall, empower ultimate survival and continuous evolution. The successful development of biofilms heavily relies on the self-production of secreted organic polymers, which may account for over 90% of the biofilm dry mass (16). These polymers contribute to the formation of the three-dimensional (3D) structure of the biofilm and facilitate interactions between cells, similar to those in multicellular eukaryotes (17). The most well-established building blocks of ECM include extracellular DNA (eDNA) (18), extracellular polysaccharides (EPS) (19), proteins (20, 21), surfactants (22), and lipids (23). Among several metals that contribute to biofilm development, calcium (Ca), the third most abundant metal in nature, has emerged as a key regulator. In eukaryotes, where Ca signaling is best studied, it is involved in almost every aspect of cellular life and death by regulating such essential processes as division, apoptosis, and immune defenses (24, 25). We recently reviewed the current understanding of the role of bacterial multicellular behavior (26) and explored the structural, signaling, and regulatory importance of Ca (27, 28). Although ample evidence supports a fundamental role of Ca in bacterial physiology, Ca signaling and regulation in bacteria remain vastly understudied.

The significant impact of Ca on biofilm formation has been reported in numerous bacterial species. Exposing cells to elevated levels of Ca triggers the switch from free-swimming to biofilm formation (26), which is mediated by its role in promoting initial adhesion to surfaces as the first step in biofilm formation (29), stabilizing and enhancing structurization of the extracellular polymeric matrix (30), as well as acting in signaling cascades that regulate the transcription of biofilm-associated mechanisms and ECM components (31–33). Ca-dependent adhesion to abiotic and cellular surfaces relies on the modification of cellular structures and macromolecules that define cell-surface properties and electrostatic interactions (34–37). These include type I and type IV pili, fimbriae (38–40), as well as teichoic acids, adhesins, lipopolysaccharide (LPS), and EPS (41). For example, Ca binds to type IV pilus-biogenesis factor, PilY1, enabling pilus extension and retraction (42). It also binds cell-surface adhesins, such as SdrC and SdrD in Staphylococcus aureus [43, 44] and LapF in *Pseudomonas. putida* (43–45), and surface proteins in *Streptococci* (46). Further, Ca strengthens biofilm matrices by forming ionic bridges between negatively charged macromolecules, such as alginate in *Pseudomonas aeruginosa* (47). Ca increases the release of and binds eDNA, another important component of the biofilm matrix, thereby increasing aggregation and contributing to biofilm formation in several Gram-positive and Gram-negative bacteria, including *S. aureus, P. aeruginosa* (48), and *Streptococcus mutans* (49).

Calcium can also generate functional crystalline biomaterials with communal functions: The formation of precisely organized calcium-based macrostructures was demonstrated robustly within *B. subtilis* biofilms and was studied extensively by us and others (50–53). These calcite scaffolds contributed to the fitness of biofilm colonies, acting as a structural scaffold supporting the 3D architecture of the colony (51, 53) and as a diffusion barrier preventing the penetration of solutes into the biofilms (52). Moreover, MicroCT X-ray analysis indicated that calcium carbonate-based 3D structures also support biofilms of the pathogens *P. aeruginosa* and *Mycobacterium abcessus* (54). The formation of controlled calcium carbonate macrostructures in phylogenetically distinct heterotrophic biofilms suggests that the function of calcium carbonate is not limited to specific species.

It has previously been demonstrated that the growth of crystalline minerals depends on the production of charged exopolymers that serve as templates in the biofilm matrix (26, 55–57). In *B. subtilis*, a master regulator, Spo0A promotes biofilm formation by enhancing the production of SinI, a small protein that acts as an antagonist to the master repressor of biofilm formation, SinR SinR directly represses two operons involved in the matrix components, namely *epsA-O,* encoding charged polysaccharides (58), and *tapA-sipW-tasA*, encoding the matrix fibrils (59–61). Both operons contribute to calcite maturation (26, 62). In addition, the repressor AbrB also represses the transcription of ECM genes [67] but has not yet been tested for biomineralization.

Despite extensive evidence that calcium promotes biofilm formation and calcite scaffold development in diverse species, it remains unclear whether biomineralization is driven simply by calcium availability and increased biomass, or whether specific calcium chemistries are required to activate matrix-dependent mineral organization (51, 63, 64). This distinction is particularly important for engineered living materials, where poorly soluble calcium-rich substrates such as calcium carbonate or construction waste are attractive feedstocks but may differ sharply from soluble organic calcium salts in bioavailability, signaling capacity, and effects on the ECM (57, 65).Separating the contributions of calcium, carbonate, and acetate is therefore essential for determining whether *B. subtilis* can directly convert exogenous mineral substrates into organized calcite structures, or whether self-generated/localized carbonate production and ECM activation are required

In this study, we compare soluble and insoluble calcium sources to decouple their effects on growth, SinI-dependent matrix regulation, colony morphology, and calcite production. We show that insoluble calcium carbonate can support growth but is insufficient to trigger robust matrix-associated biomineralization in wild-type *B. subtilis*, whereas ECM-overproducing mutants partially overcome this barrier.

## Materials and Methods

### Strains

All *B. subtilis* strains utilized in this study are derivatives of the undomesticated wild-type strain NCIB 3610, characterized by its ability to form robust, complex biofilms. To dissect the regulatory pathways linking mineralization to biofilm formation, we employed specific isogenic mutants, including the matrix-overproducing strains (Δ*sinR* and Δ*abrB* deletion mutants) and the P*sinI*-lux transcriptional reporter strain for real-time monitoring of matrix gene expression. A comprehensive list of all strains, including their specific genotypes and sources, is provided in Supplementary Table S1.

### Growth Conditions and Imaging

Bacterial cultures were prepared by inoculating a single colony, isolated on lysogeny broth (LB) agar (BD Difco™, Thermo Fisher Scientific, Waltham, MA, USA), into 3 mL of LB broth. The cultures were incubated at 30 °C with agitation for 4 hours to reach the mid-logarithmic phase. To initiate biofilm formation, 2 µL aliquots of the bacterial suspension were spotted onto solid B4 biomineralization medium (composition: 0.4% yeast extract, 0.5% glucose, and 1.5% agar) supplemented with Calcium Acetate (Ca (CH_3_COO)_2_) at final concentrations of 2, 10, or 20 mM. Control groups were cultivated on un-supplemented B4 medium. To evaluate substrate specificity and distinguish the effects of the calcium cation from the counter-anions, comparative assays were conducted on modified B4 media supplemented with Calcium Carbonate (CaCO_3_; 0.2, 1, or 2 mg/mL, corresponding approximately to 2, 10, or 20 mM calcium), Sodium Acetate (NaCH_3_COO; 2, 10, or 20 mM), or Sodium Carbonate (Na_2_CO_3_; 2, 10, or 20 mM). All plates were incubated at 30 °C for up to 24 days. Macroscopic colony morphology was documented using a Nikon Coolpix P950 digital camera.

### Calcium carbonate crystal Extraction

To produce crystals, strains were streaked on biomineralization-promoting solid medium and incubated at 30 °C for up to 24 days. Crystals were visible after 18 days of incubation. Crystals were collected as previously described by us in Oppenheimer-Shaanan et al. (53) With some modifications, agar samples were slightly bleached with 6% sodium hypochlorite for 1 min to remove organic matter, then washed twice with DDW and dehydrated in acetone.

### FTIR Spectroscopy

FTIR spectra of the produced crystals were acquired by using a NICOLET Summit X FTIR spectrometer (Thermo Scientific, Pittsburgh, PA, USA). A few milligrams of the sample were used for each measurement. Infrared spectra were obtained at 4 cm−1 resolution for 32 scans. The baselines for the height measurements of the v2, v3, and v4 peaks were determined as previously done by Oppenheimer-Shaanan et al (53). The v2, v3 and v4 heights were normalized to a v3 height of 1,000, corresponding to 1.0 a.u.65

### Kinetic Growth and Expression Assays

Kinetic measurements were performed using the *B. subtilis* transcriptional reporter strain IKbs45056 (*amyE*::P*sinI*-lux). Pre-cultures were grown in LB broth at 30 °C with orbital shaking at 200 rpm for 4 hours to reach the mid-logarithmic phase. Subsequently, cultures were diluted at a 1:100 ratio into fresh liquid B4 medium supplemented with Calcium Acetate (Ca(CH_3_COO)_2_), Sodium Acetate (NaCH_3_COO), or Sodium Carbonate (Na_2_CO_3_) at final concentrations of 2, 10, or 20 mM and Calcium Carbonate (CaCO_3_), at final concentrations of 0.2, 1 or 2 mg/mL. The suspensions were distributed into a 96-well white polystyrene microplate with a clear bottom (Visiplate, Wallac). The microplate was incubated in a Tecan Infinite 200 Pro multimode reader (Tecan, Männedorf, Switzerland) at 30 °C for 24 hours. Bacterial growth (𝑂𝐷_600_) and luciferase-derived luminescence (sensitivity setting: 200) were monitored at 10-minute intervals. Data processing, including background subtraction, averaging of quadruplicate biological replicates, and signal normalization, was executed automatically using the MatTek computational package.

### Computational Data Analysis

Rigorously testing hypotheses about microbial responses requires dissecting the precise temporal coupling among bacterial growth, gene expression, and mineralization dynamics. This presents a significant computational challenge, as high-throughput kinetic assays are often plagued by signal noise, non-linear artifacts, and well-to-well variability, which can obscure subtle physiological patterns [71,72]. To address this methodological gap and enable precise quantification of biomineralization competence, we developed “MatTek,” an automated computational framework implemented in MATLAB. This platform integrates connectivity-based segmentation, automated baseline alignment, and robust sliding-window algorithms to extract high-resolution metabolic parameters from complex kinetic data, thereby addressing limitations of traditional manual analyses [73]. The computational parsing module within the developed MatTek software was originally optimized for the data layout produced by Tecan Spark/2.0 readers. However, because the software uses a template-based, instrument-agnostic framework, it seamlessly parses the standardized ASCII/Excel output files generated by the actual acquisition hardware, the Tecan Infinite 200 Pro.

This dedicated computational platform is implemented within the MATLAB R202X environment (MathWorks Inc.). Optimized for the high-throughput output of Tecan 2.0 microplate readers, this in-house pipeline offers extended feature-extraction capabilities compared to established tools such as GrowthRates 3.0 [74,75] and our previously reported GROOT software [76]. The analysis framework was structured into distinct algorithmic modules that executed end-to-end data curation, advanced signal processing, kinetic modeling, and statistical evaluation. A key feature is its interactive graphical interface that allows users to selectively assign wells from a microplate to different experimental groups, with real-time visual feedback and persistent selection tools. The software then generates publication-ready visualizations, including growth curves with error bands, 2D scatter plots correlating luciferase expression with optical density, (OD) and 3D trajectory plots showing temporal dynamics.

### Automated Data Curation and Segmentation

To ensure high-throughput data integrity, raw export files from the microplate reader were processed via an automated parsing algorithm. A connectivity analysis algorithm (bwconncomp) was employed to scan the raw data matrix and identify contiguous numeric clusters, allowing for the automated segregation of OD_600_ and Luminescence (𝐿𝑢𝑚) datasets. This process effectively filtered out metadata and non-numeric artifacts without manual intervention. Following segmentation, datasets were computationally screened for anomalies; non-finite values were excluded, and negative values resulting from background subtraction were clamped to zero using a rectification function, 𝑓(𝑥) = max(0, 𝑥), to maintain physical validity.

### Signal Pre-processing and Baseline Alignment

To eliminate experimental artifacts arising from well-to-well variability, such as initial optical offsets or meniscus effects, a Baseline Alignment Algorithm was applied before kinetic analysis. The initial OD of the control group at time *t* = 0, denoted as *OD*_*ctrl*_(*t*_0_), was established as the global zero-reference. The corrected growth trajectory for each sample *i*, denoted as *OD*^′^(*t*), was calculated by applying a scalar shift to align its starting point with this reference, as shown in Eq. (1):

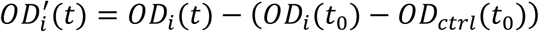

Subsequently, gene expression signals were normalized to bacterial biomass to yield Specific Luminescence Activity (𝑅𝐿𝑈/*OD*) via the ratio described in Eq. (2):

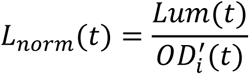

### Bacterial Growth Kinetics

Quantitative growth parameters were extracted from the processed *OD*_600_ curves using regression models. The Specific Growth Rate (𝜇) was calculated as the slope of the natural logarithm of biomass during the exponential phase (*t* ∈ [*t*_*start*_, *t_end_*]), utilizing Ordinary Least Squares (OLS) regression (Eq. 3):

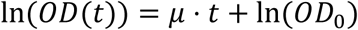

Goodness-of-fit was assessed via the Coefficient of Determination (*R*^2^). Additionally, Lag Time (𝜆) was formally defined as the minimal time required for the culture to exceed a predefined growth threshold (𝛿 = 0.1) relative to its baseline (Eq. 4):

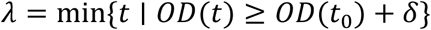

Total bacterial growth was quantified by calculating the Area Under the Curve via numerical integration using the Trapezoidal Rule, approximated by Eq. (5):

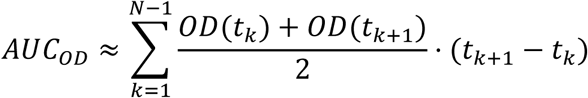

### Reporter Expression Dynamics

Luciferase expression kinetics were characterized using a robust Sliding Window Algorithm. The Maximal Expression Rate (*V*_*max*_) was determined by scanning the kinetic profile using a moving time-window of width *W*. For each window ending at time *t*_*k*_, the local expression rate was estimated via linear regression (𝛽_1_), and the maximal rate was defined as the global maximum of these local slopes (Eq. 6):

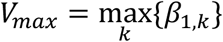

Expression Efficiency (𝜂) was calculated to describe the system’s responsiveness, defined as the ratio of peak luminescence intensity to the time elapsed from induction to saturation (Eq. 7):

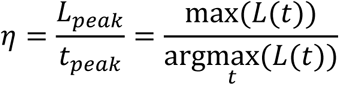

The Expression Duration (Δ*t_exp_*) was calculated by defining a temporal set 𝒯_*active*_where the signal exceeded a dynamic threshold (𝜃) set at 10% of the peak intensity (Eq. 8); the duration was defined as the span of this set (Eq. 9):

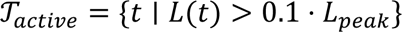

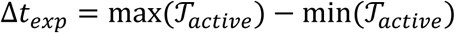

Finally, Fold Change (*FC*) was quantified by normalizing the peak intensity of the sample against the reference control group (Eq. 10):

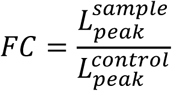

### Multi-dimensional Visualization and Statistical Analysis

To visualize the dynamic coupling between growth and expression, data were mapped into a 3D coordinate system defined by the vector 𝑣⃗(*t*) = [*OD*(*t*), 𝐿𝑢𝑚(*t*), *t*], illustrating the state evolution over time. Data aggregation involved computing the arithmetic mean and standard deviation (SD) across 𝑛 replicates (𝑀𝑒𝑎𝑛 ± 𝑆𝐷). Significance levels were visualized using 𝑝-value heatmaps, with statistical significance set at 𝛼 < 0.05.

### Amorphous Calcium Carbonate Preparation (ACC)

ACC was prepared using a chilled precipitation method(66). Briefly, Solution A (0.424 g sodium carbonate in 20 mL of 2M NaOH, diluted to 200 mL with deionized water) and Solution B (0.588 g calcium chloride in 200 mL deionized water) were cooled in an ice bath. Solution A was then rapidly poured into Solution B under vigorous stirring to induce immediate precipitation to achieve a controlled, equimolar precipitation of transient ACC. The mixture was promptly subjected to vacuum filtration, and the obtained precipitate was washed several times with co(66)ld acetone to halt crystallization. The final powder was dried in vacuo overnight at room temperature.

### Bacterial Culture Preparation and Bio-cementation Assay

#### Inoculum Preparation

Bacterial strains were revived from frozen stocks by streaking onto LB agar plates, followed by incubation at 30 °C for 24 hours. A single isolated colony was then inoculated into 2 mL of LB supplemented with selective antibiotics and incubated for an additional 24 hours at 30 °C. To initiate the biomineralization phase, the entire 2 mL pre-culture was transferred into 20 mL of B4-Ca medium supplemented with specific antibiotics to maintain plasmid or strain selection. Antibiotic concentrations were applied as follows: for the control strain (wild type carrying antibiotic resistance genes, the medium was supplemented with erythromycin and Chloramphenicol (to a final concentration of 100 ng/ml and 10 µg/ml, respectively); for the Δ*abrB* mutant, Kanamycin was added to a final concentration of 10 µg/ml); and for the Δ*sinR* mutant, Spectinomycin was added (100 µg/ml).

### Flow cytometry analysis of expression from P*sinI*-GFP

Flow cytometry was used to quantify activation of the *sinI* promoter at the single-cell level. Wild-type *B. subtilis* and the *amyE*::P*sinI*-GFP reporter strain were grown on solid B4 medium with or without calcium supplementation. Briefly, 2 µL aliquots of bacterial culture were spotted onto B4 agar plates containing no treatment, calcium acetate, calcium carbonate, or calcium chloride at 0.25% w/v. Plates were incubated at 30 °C for 24 h. Cells were then recovered from the agar surface, fixed with 4% paraformaldehyde, and analyzed using by CytoFLEX S flow cytometer. For each sample, 10,000 events were collected. The GFP-positive population was defined relative to the wild-type/background control, and representative plots were selected from the reporter replicates closest to the mean GFP-positive fraction for each condition.

#### Stereomicroscopy of colony morphology

Colony morphology was examined using a Leica stereomicroscope equipped with a PLANAPO 1.0× objective, a FLUOIII module, a Leica TL3000 Ergo transmitted-light base, and an attached Leica K3M camera. Wild-type *B. subtilis* cultures were prepared as described above, and 2 µL aliquots were spotted onto B4 agar plates without supplementation or supplemented with 10 mM calcium acetate or 10 mM calcium chloride. Plates were incubated at 30 °C for 11 days. Representative colonies were imaged from three independent plates per condition. Overview images were acquired at 0.8× magnification, and higher-magnification images were acquired at 5× to examine colony surface morphology and crystal-associated structures.

#### Construction Waste Consolidation

To evaluate the macroscopic consolidation capability of the bacterial cultures, a molding assay was performed using ground construction waste as an aggregate scaffold. Custom silicone molds were fabricated via 3D printing and packed uniformly with the waste substrate. A 20 mL aliquot of the prepared bacterial suspension in B4-Ca medium was introduced onto the matrices until saturation, and excess liquid was removed via aspiration. The inoculated molds were incubated at 30°C for 48 hours to facilitate microbially induced calcium carbonate precipitation (MICP) and subsequent consolidation.

Following incubation, the solidified structures were demolded, and their structural integrity was qualitatively assessed upon handling. Pore distribution within the consolidated matrices were subsequently characterized by digital image analysis.

## Results

### The effect of Calcium sources on Biofilm formation, pigmentation, and colony structure

In a saturated solution (e.g., To assess the possible roles of biomineralization in biofilm development, we grew wild-type *B. subtilis* cells in media containing calcium acetate or calcium carbonate as a calcium source. In un-supplemented B4 medium, colonies remained flat and translucent white, with no complex morphology (Fig. 1a and Fig. S1). By contrast, calcium acetate altered colony architecture in a concentration-dependent manner, producing pigmented and structured colonies with visible mineral-associated zones, most prominently at 10 and 20 mM (Fig. 1a). A magnified view of a calcium-acetate-grown colony highlights the mineral-associated colony edge used for downstream crystal analysis (Fig. 1b). Next, Fourier transform infrared (FTIR) spectroscopy was used to identify the precipitated mineral observed within biofilm exposed to calcium acetate and to distinguish between different crystalline polymorphs of CaCO_3_: calcite, aragonite, and vaterite. We removed the organic matter by a hypochlorite priming step, and analysis was performed on the inorganic matter only. The infrared calcite spectrum exhibits three characteristic peaks in the region of 400–4000 cm−1, designated as v_2_, v_3_, and v_4_ (67). Calcium carbonate ν_3_ peak asymmetric stretching band is typically expected near 1425 cm−1 for calcite, 1490 cm−1 for vaterite, and 1475 cm−1 for aragonite; however, these peaks can shift down in the presence of lattice defects or metal impurities (68). The calcium carbonate ν_2_ peak is expected at 875 cm−1 for calcite, 850 cm−1 for vaterite, and 855 cm−1 for aragonite. Finally, the calcium carbonate ν_4_ peak is expected at 713 cm−1 for calcite, 750 cm−1 for vaterite, and 715 cm−1 for aragonite [52, 53]. The FTIR spectra of the putative calcium carbonate minerals that were collected from the edges of the biofilm were typical of calcite (Figure 1c).

**Figure 1.**
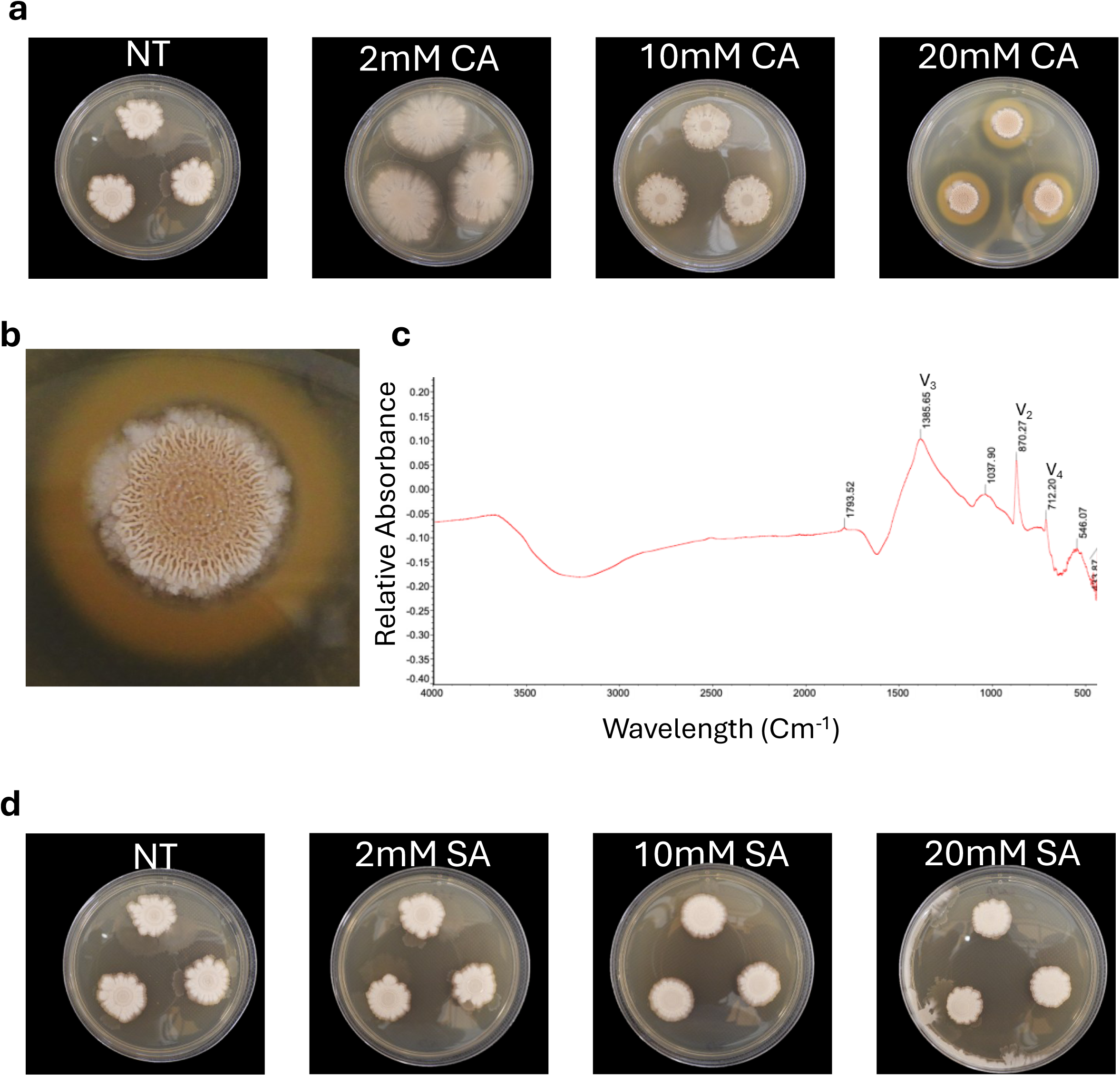
Calcium-driven mineralization in B4 medium. Representative images of *B. subtilis* colonies grown on B4 biomineralization medium without treatment (NT) or supplemented with calcium acetate (CA; 2, 10, or 20 mM). (b) Magnified view of a calcium-acetate-grown colony showing the structured, pigmented mineral-associated colony edge. (c) Fourier-transform infrared (FTIR) spectrum of crystals collected from biofilm edges, consistent with calcite based on the ν_2_, ν_3_, and ν_4_ absorption bands. (d) Representative colonies grown on B4 medium supplemented with sodium acetate (SA; 2, 10, or 20 mM), showing that acetate alone does not reproduce the calcium-acetate-associated mineralized colony morphology. Results represent one out of three independent experiments, performed with five technical repeats. Control of the same batch is used for SA and CA with source data deposited here: <u>folder</u>.

To rule out the possibility that biomineralization of detectable calcite crystals is induced by acetate rather than by calcium, we repeated the experiment using equimolar concentrations of sodium acetate. Sodium acetate did not induce 3D pattern formation or crystalline mineral accumulation within the colony (Fig. 1d). Similarly, sodium carbonate did not induce the calcium-acetate-associated colony morphology (Fig. S2).

Interestingly, while amorphous calcium carbonate (an intermediate state commonly stabilized in biological systems) could induce moderate crystal precipitation (borderline to detection limit), it was active only at concentrations above 1 mg/mL (Fig. S3). This observation highlights the thermodynamic bottleneck that defines the transition from metastable phases to stable mineral structures. In many biological systems, Amorphous Calcium Carbonate (ACC) is the preferred Calcite precursor. However, because it lacks long-range crystalline order, its transformation into a stable mineral such as calcite or aragonite requires overcoming a specific energy barrier, which is provided here by the bacterial cells.

### Calcium Supports Cellular Growth and Enhances Carrying Capacity

Our results indicated that bacteria generate crystalline calcium carbonate, which could, by itself, affect microbial growth. To accurately assess the effects of calcium, carbonate, and acetate on growth, we measured cell turbidity over time under each condition. In the biomineralization medium, both calcium acetate and calcium carbonate influenced microbial growth significantly (Fig. 2a and b). As crystalline material was observed only at high calcium acetate concentrations (>= 10 mM), this result could indicate a non-linear relationship between biomineralization and growth, and that biomineralization is not a readout of bacterial biomass per se. Interestingly, saturated calcium carbonate solutions (equivalent to 20 mM) significantly promoted microbial growth, yet no crystalline material was observed within the colony (Fig. 2c). Furthermore, sodium hydroxide lysis treatment yielded no quantifiable inorganic bleach-insoluble mineral even 20 days post-inoculation (Table S2).

**Figure 2.**
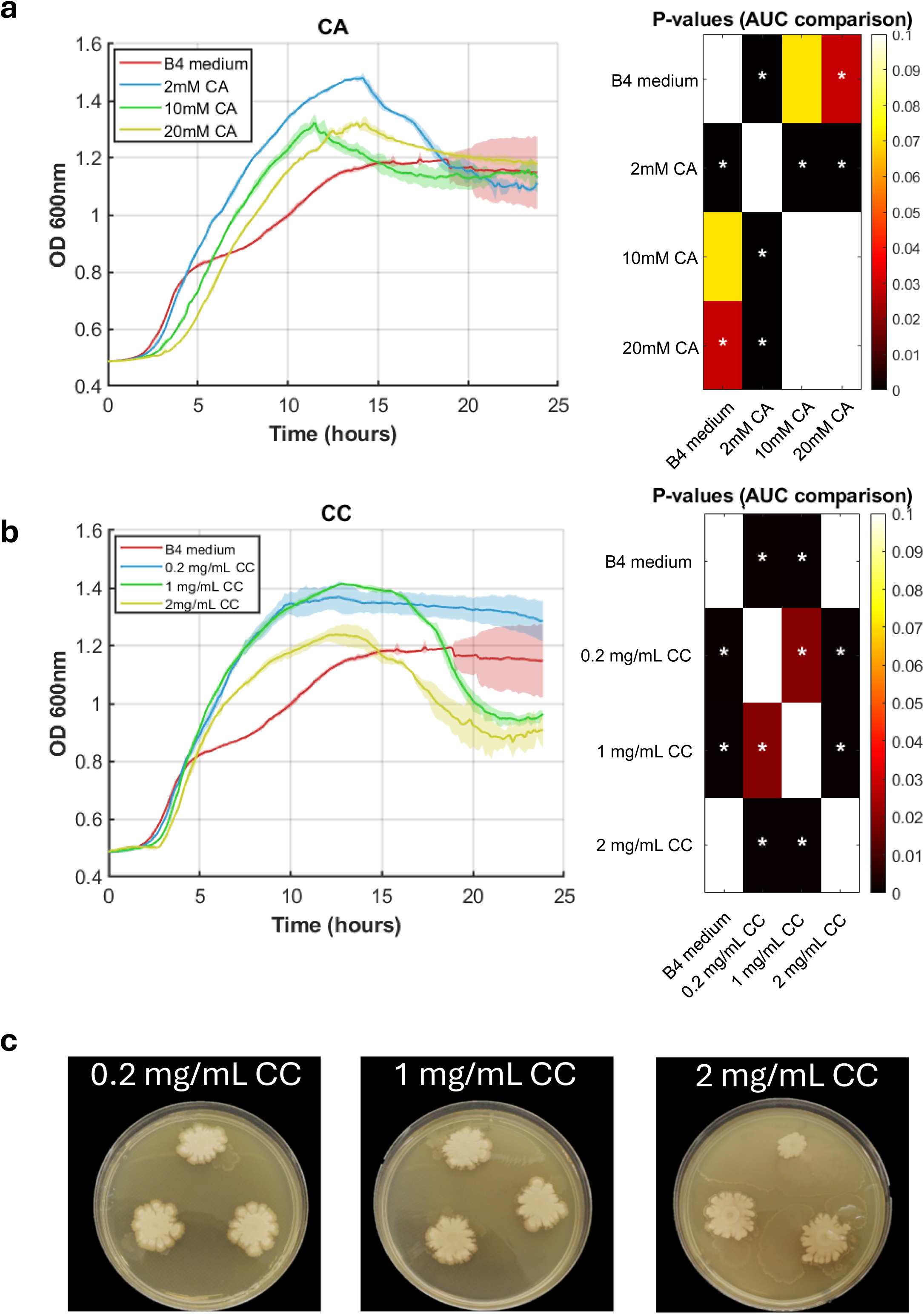
Growth dynamics across different calcium salt concentrations. Growth kinetics monitored by OD_600_ over 24 h in liquid B4 medium supplemented with (a) calcium acetate (CA; 2, 10, or 20 mM) or (b) insoluble calcium carbonate (CC; 0.2, 1, or 2 mg/mL, corresponding approximately to 2, 10, or 20 mM calcium). B4 medium without added calcium served as the control. Shaded regions represent the standard deviation. Heatmaps show P values from area-under-the-curve (AUC) comparisons among conditions. (c) Representative colonies grown on solid B4 medium supplemented with CC, showing that CC supports growth-associated colony formation without inducing visible calcite accumulation or the calcium-acetate-associated structured morphology. Results represent one out of three independent experiments, performed with five technical repeats.

Utilizing the MatTek platform (materials and methods), we analyzed microbial growth in detail. The platform provides a complete workflow for handling OD and luminescence (luciferase) measurements commonly obtained from bacterial growth and gene expression studies. The system begins with flexible data preparation, automatically detecting and extracting multiple data blocks from Excel spreadsheets, then proceeds through progressive analysis stages including raw data visualization, individual group comparisons, and advanced statistical evaluations. Beyond visualization, MatTek incorporates analytical modules that compute metrics such as growth rates, lag times, peak expression values, and area-under-curve measurements, as well as statistical comparisons between groups using ANOVA and t-tests. The modular architecture enables seamless integration of these stages of analysis while maintaining data integrity throughout the pipeline, making it suitable for systematic analysis of complex biological experiments involving multiple treatment groups and time-series measurements.

First, we quantified growth kinetics—specifically lag phase duration, growth rate, and carrying capacity—across a range of salt concentrations (Fig. 3a). Calcium acetate enhanced the growth rate at 2, 10, and 20 mM; however, the growth induction was more pronounced at 2 mM than under biomineralization-inducing conditions (10 and 20 mM). While salt addition elongated the lag phase, the overall effect was not substantial enough to significantly alter the doubling time relative to the control (Fig. 3a). In contrast, sodium acetate failed to induce carrying capacity (maximal OD) significantly and had a mild effect on the growth rate (Fig. 3b). These results indicate that calcium is an underappreciated essential metal in the relatively rich biomineralization medium (53) and that carbonate buffering further supports microbial growth. To confirm these findings and rule out the possibility that the observed differences in optical density were due to extracellular polymeric substance (EPS) secretion rather than true cell proliferation, we quantified colony-forming units (CFUs). Crucially, all assays (Figs. 2-4, S5 and S6) were performed under continuous shaking in non-biofilm conditions. This analysis confirmed that calcium acetate indeed enhances true cell growth, a phenotype that was fully recapitulated in the matrix-deficient mutant background (Fig. S4).

**Figure 3.**
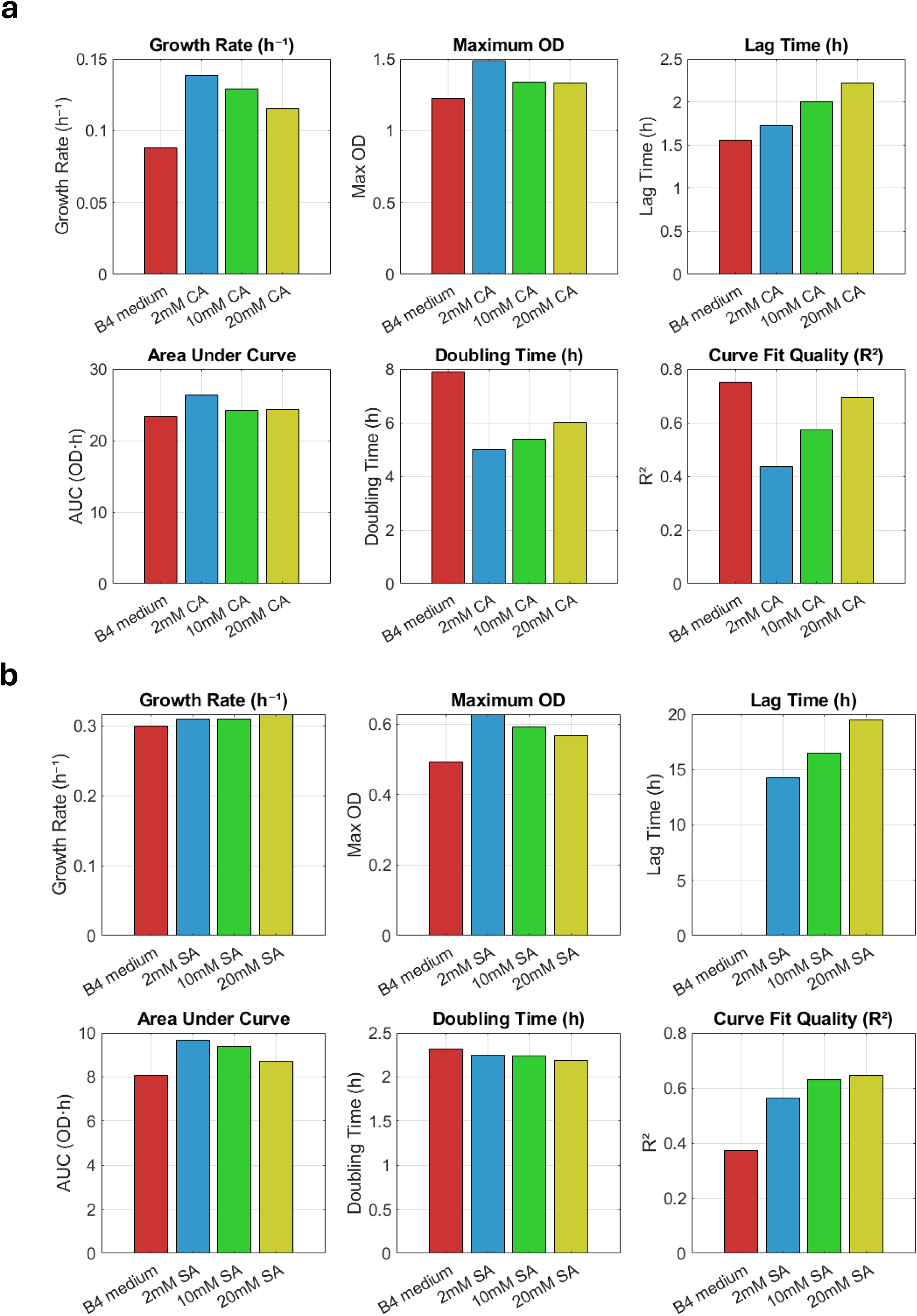
Automated kinetic analysis of growth in response to acetate salts. Quantitative growth parameters extracted using the MatTek platform, including growth rate, maximum optical density (Max OD), lag time, area under the curve (AUC), doubling time, and curve-fit quality (R2). Panels (a) and (b) correspond to calcium acetate and sodium acetate, respectively. Results represent one out of three independent experiments, performed with five technical repeats.

**Figure 4.**
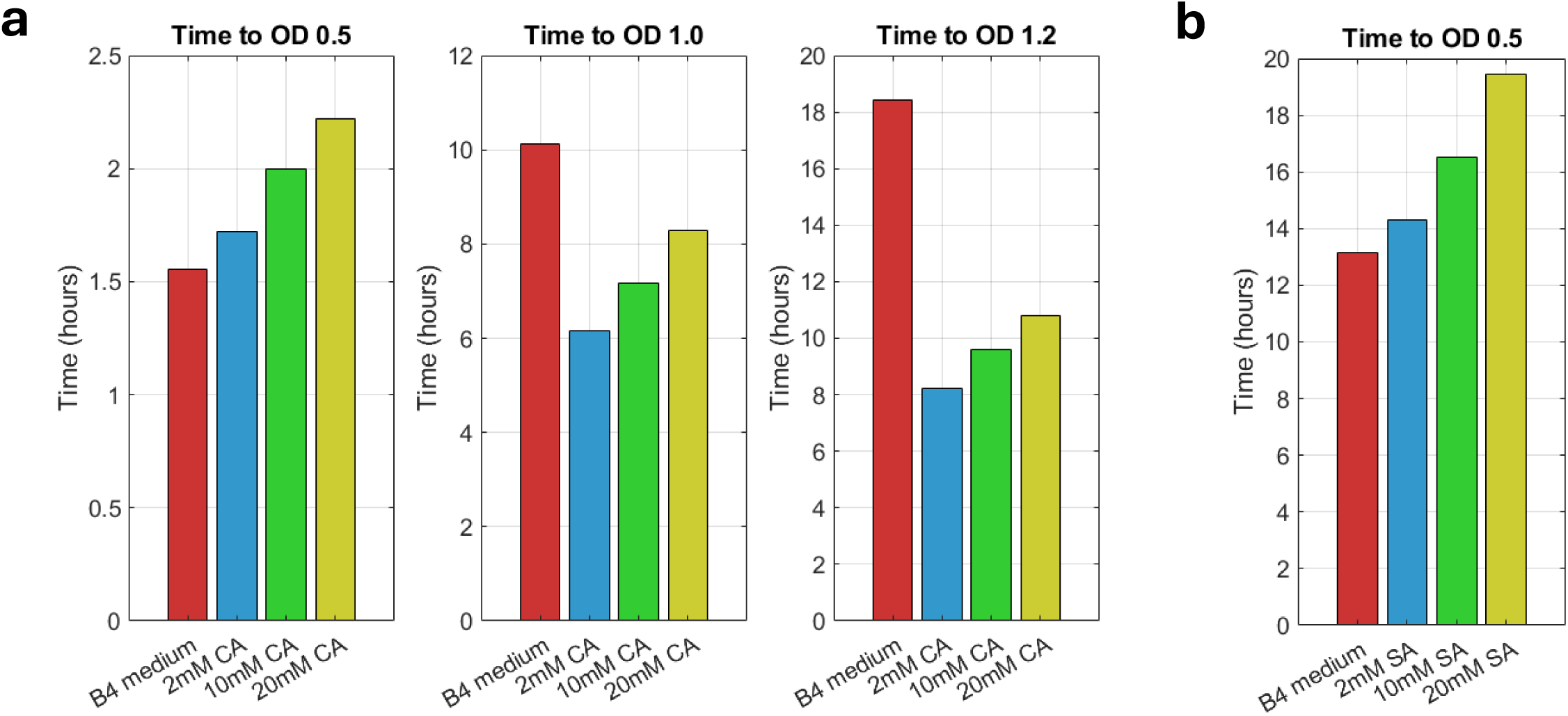
Acetate does not support enhanced carrying capacity. Time-to-threshold analysis was used to determine the time required for cultures to reach defined OD_600_ values. (a) Calcium acetate conditions were compared with B4 medium at thresholds of OD_600_ 0.5, 1.0, and 1.2. (b) Sodium acetate conditions were compared with B4 medium at OD_600_ 0.5, consistent with the lower carrying capacity observed under sodium acetate treatment. performed with at least three technical repeats.

To evaluate differential contributions, we analyzed the lag phase duration, which showed a significant reduction at 20 mM (Fig. 4a and b). Both calcium acetate and sodium acetate dramatically shortened the lag period; rapid quantification of the time-to-threshold OD using MatTek software facilitated these streamlined comparisons. Strikingly, despite its poor solubility, insoluble calcium carbonate contributed similarly to growth kinetics according to this parameter as calcium carbonate (Fig. S5), further suggesting effective bacterial utilization of the feedstock crystalline powder.

Given that calcium carbonate was not inert to microbial growth, we investigated the distinct roles of soluble sodium carbonate and insoluble calcium carbonate. To our surprise, calcium carbonate supported growth at levels comparable to those of soluble calcium acetate (Fig. 5a), whereas sodium carbonate had a negligible impact (Fig. 5b). Notably, neither carbonate source triggered characteristic colony morphogenesis nor calcite precipitation within the biofilm (Figs. 2c and Table S2). These results further decouple microbial growth from the physiological parameters required for biomineralization and structural maturation, and carbonate has little or no effect on triggering these changes when applied in isolation. Additionally, as sodium carbonate and calcium carbonate induce similar pH shifts, this observation suggests that the growth regulation is independent of alkalinity.

**Figure 5.**
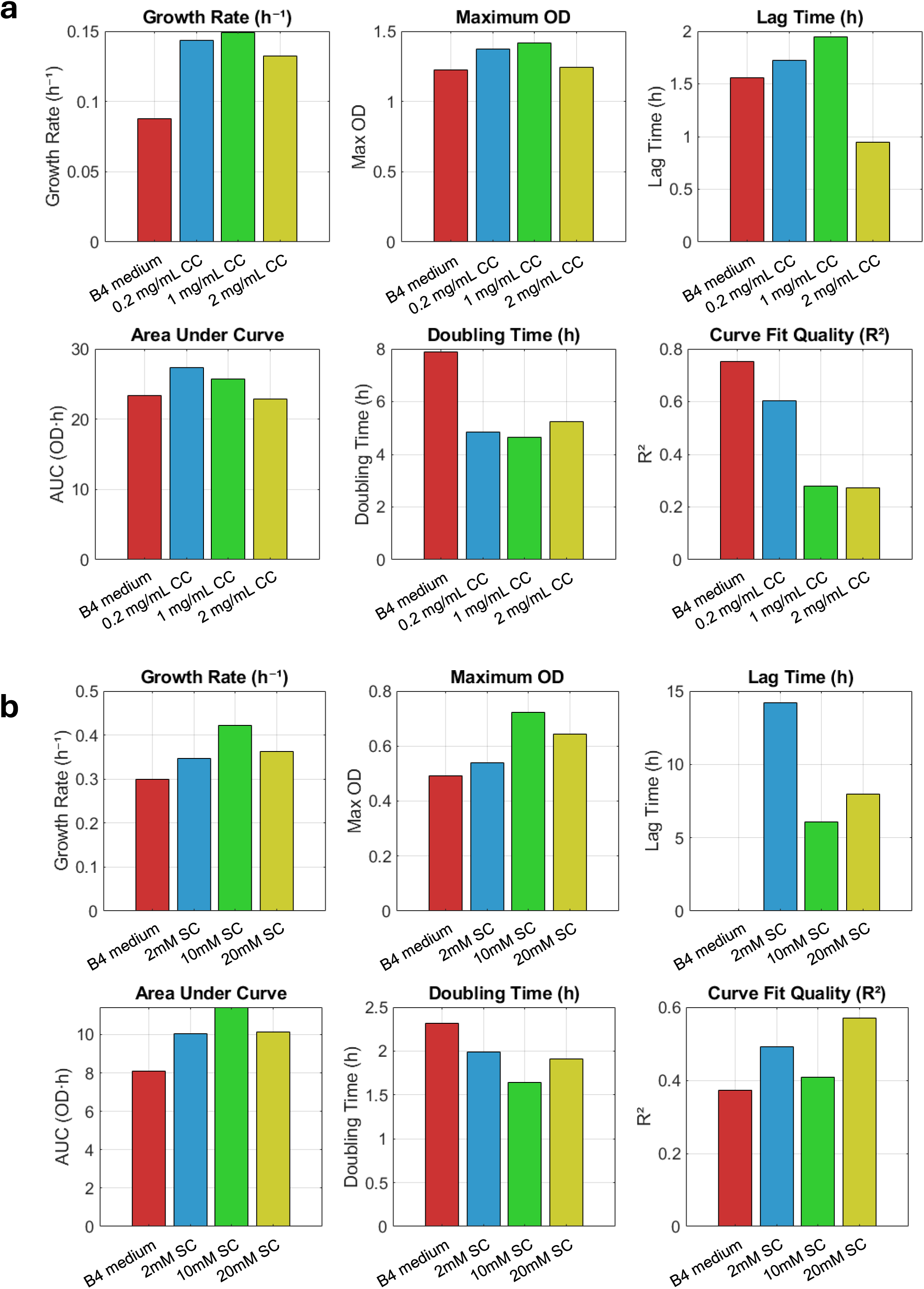
Calcium from Insoluble Calcium Carbonate supports enhanced carrying capacity (maximal OD) and reduced doubling time. Quantitative growth repeats. extracted using the MatTek platform, including growth rate, Max OD, lag time, area under the curve (AUC), doubling time, and curve-fit quality (R2). Panels (a) and (b) correspond to calcium carbonate and sodium carbonate, respectively. Results represent one out of three independent experiments, performed with at least three technical repeats.

### The Expression from the SinI Promoter is calcium-dependent

The protein SinI acts as an antagonist to SinR, the master repressor of biofilm formation [66, 67][68]. Consequently, *sinI* transcription serves as a robust proxy for ECM production and, by extension, the biomineralization potential of *B. subtilis*. Using our automated kinetic framework, we analyzed the luminescence of a P*sinI*-lux transcriptional reporter across a range of concentrations of calcium acetate, calcium carbonate, sodium acetate, and sodium carbonate (Fig. 6a-e). To account for differences in biomass, expression was normalized to growth (RLU/OD), thereby isolating the per-cell expression efficiency.

**Figure 6.**
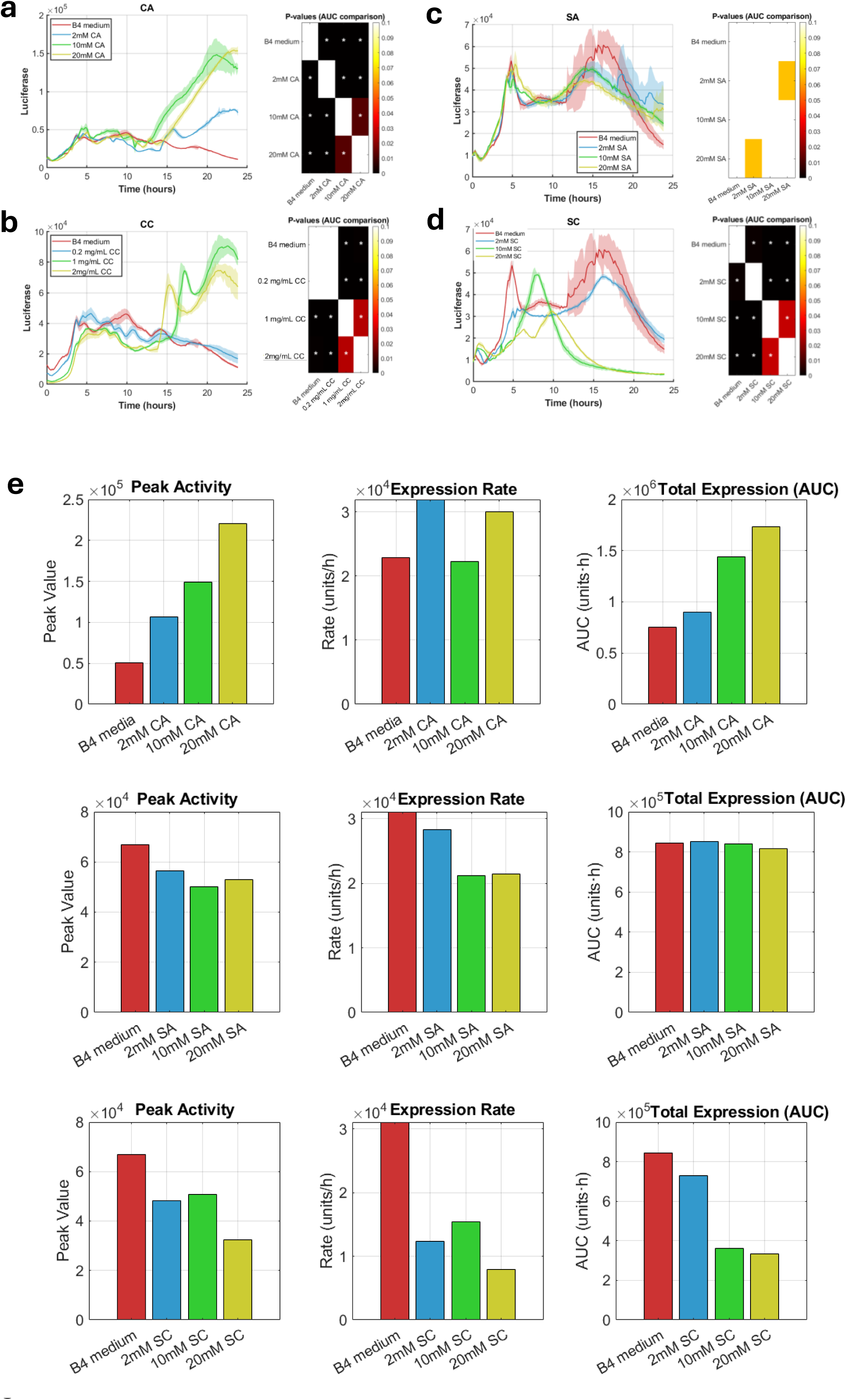
Calcium increases both the duration and intensity of SinI expression. Temporal luminescence profiles of the P*sinI*-lux transcriptional reporter during growth in B4 medium supplemented with (a) calcium acetate (CA; 2, 10, or 20 mM) or (b) carbonate (CC; 0.2, 1, or 2 mg/mL), (c) sodium acetate (SA) or (d) sodium carbonate (SC) (2, 10, or 20 mM). (e) Quantitative reporter-expression metrics derived using MatTek are shown for calcium acetate, sodium acetate, and sodium carbonate, including peak activity, expression rate, and total expression (AUC). Results represent one out of three independent experiments, performed with at least three technical repeats.

The SinI expression profile exhibited a distinct bimodal distribution, comprising early and late expression zones, in the presence of calcium sources, whereas it remained unimodal in untreated controls (Fig. 6a and b). Although both calcium acetate and calcium carbonate induced total expression at high concentrations (Fig. 6a and b), their impacts on expression efficiency differed significantly their impacts on expression efficiency differed significantly (e.g., the total luminescence values at peak reaction were distinct).

While calcium carbonate supported growth effectively (suggesting active solubilization), its poor aqueous solubility likely limits the steady-state concentration of available intracellular calcium. This bioavailability constraint may explain why calcium carbonate yielded the lowest expression efficiency among the tested calcium sources, despite triggering a high initial expression rate (Fig. 6b). Neither sodium or acetate were efficient inducer of transcription from *sinI* promoter (Fig. 6c). Moreover, carbonate downregulated reporter output when provided as a sodium salt (Fig. 6d). Since luciferase is a short-lived reporter, this analysis provides high temporal resolution and underscores that calcium—rather than the acetate or carbonate counter-ions—is the primary driver of the observed shifts in expression values (Fig. 6e).

To extend this analysis to counter-ion effects, we quantified the same reporter-derived parameters for sodium acetate and sodium carbonate; sodium acetate produced only modest changes relative to B4 medium, whereas sodium carbonate reduced peak activity, expression rate, total expression, and fold change (Fig. S6).

We did not perform a comparative extraction of expression parameters for calcium carbonate and calcium acetate because their bioavailability profiles differ significantly. However, the finding that *B. subtilis* can utilize high concentrations of CaCO_3_ to support growth—but not to induce *sinI* expression (Fig. 5a and Fig. 6b)—suggests that these two physiological parameters are functionally decoupled. The lack of robust biomineralization in the presence of CaCO_3_ warrants further investigation into the specific signaling thresholds required for mineral nucleation. Notably, flow cytometry analysis (Fig. S7) confirmed a robust induction of SinI expression, with a distinct advantage observed for calcium acetate over both calcium carbonate and calcium chloride. This finding effectively decouples SinI expression from calcium solubility.

### Physiological Decoupling of Matrix Gene Expression with Biomineralization

Our findings support a model in which *B. subtilis* coordinates mineral morphology with ECM secretion as an adaptive strategy to optimize metabolic longevity and productivity within the biofilm microenvironment. To address the observed lack of synchronization between calcite nucleation and matrix-driven growth—particularly under calcium carbonate (CC) conditions—we used two hyper-biofilm-forming mutants, *ΔsinR* and *ΔabrB,* that overexpress the ECM. Macroscopic analysis indicated that the deletion of *sinR* enabled the formation of mineralized wrinkles, while the *ΔabrB*, background resulted in a significantly thicker colony phenotype on calcium carbonate (Fig. 7a).

**Figure 7.**
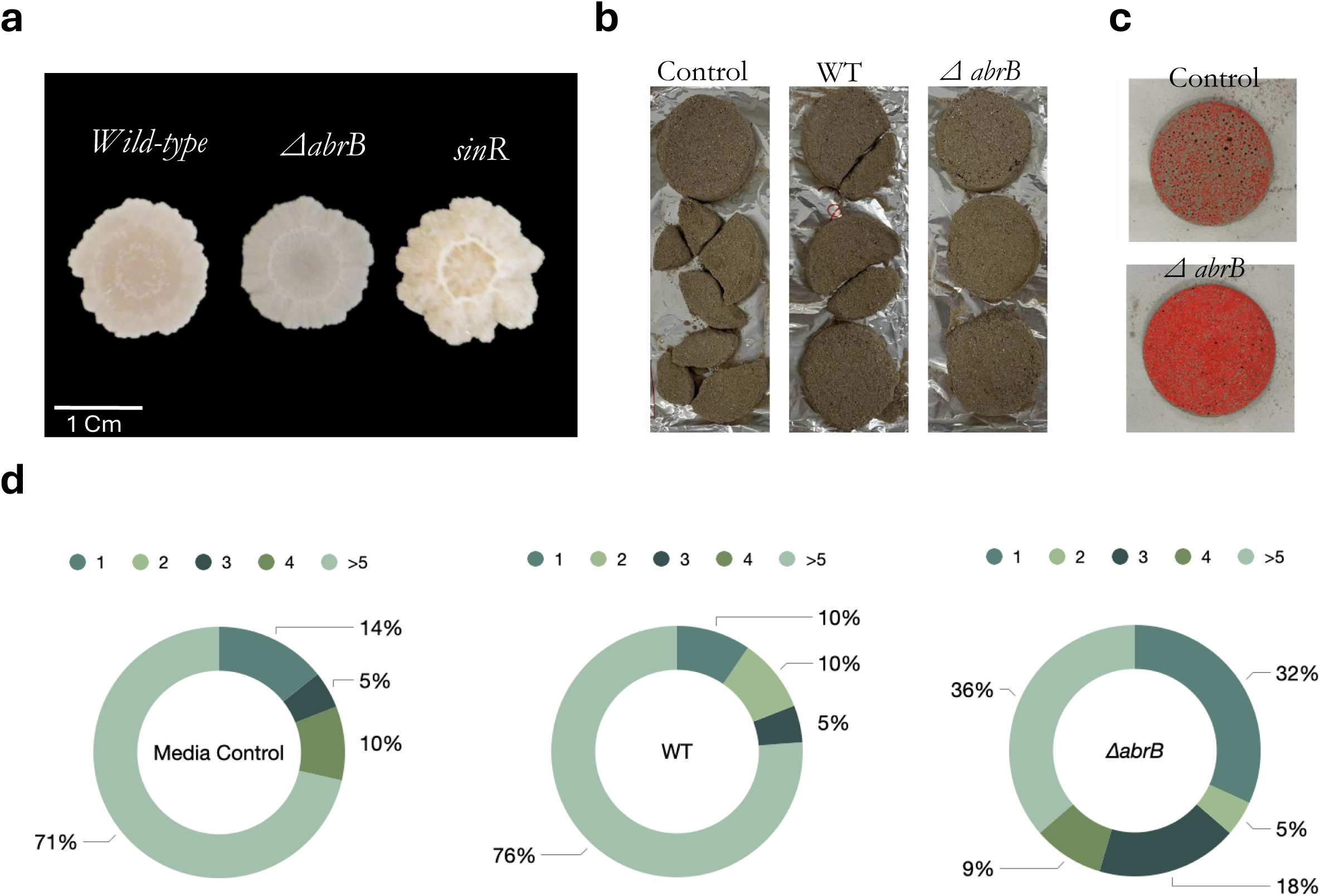
ECM overproduction partially overcomes barriers to calcium-carbonate-based consolidation. (a) Representative macroscopic colony images of wild-type, Δ*abrB*, and Δ*sinR B. subtilis* strains grown under calcium-carbonate conditions, showing strain-dependent differences in colony morphology. Results represent one out of three independent experiments, performed with at least three technical repeats. (b) Construction-waste aggregates treated with control medium, wild-type cells, or Δ*abrB* cells. (c) Image-analysis overlay of control and Δ*abrB*-treated aggregates. The Δ*abrB* mutant produced more cohesive consolidated discs than the control or wild-type strain, consistent with enhanced matrix-mediated aggregate stabilization. (d) Disk aggregation parameters across multiple experiments (7 independent repeats, performed in triplicate).

Therefore, the comparative efficacy of *B. subtilis* strains in the bio-consolidation of grinded construction waste (Figs. 7b and c). While the control and wild-type (WT) samples exhibited significant failure to consolidate and fragmentation, the *ΔabrB* mutant had significantly improved performance in consolidating the waste into cohesive, intact discs. This superior performance of the *ΔabrB* strain suggests that its hyperproduction of ECM provides the necessary biological adhesive materials to bridge and stabilize heterogeneous construction waste particles (Fig. 7d). By effectively templating calcium carbonate precipitation within its dense matrix, the *ΔabrB* mutant creates a robust mineralized network that enhances the mechanical integrity of the composite, highlighting its potential for sustainable industrial applications in waste valorization and bio-cementation.

## Discussion

Biomineralization *B. subtilis* has been extensively studied as a promising route for developing engineered living materials (55, 64). Previous studies have consistently demonstrated that the efficiency of biomineralization is highly dependent on the calcium source. Organic calcium salts, such as calcium acetate and calcium lactate, yield significantly higher bioconversion efficiencies than inorganic sources such as calcium nitrate or chloride. This advantage has been largely attributed to the metabolic utilization of the organic anion as a carbon source, which promotes local alkalinity (69)-although alkalinity per se was found by us as insufficient for biomineralization (Fig. 1, (53)).

This observed metabolic advantage of organic carbon feedstocks aligns closely with established calcium carbonate precipitation paradigms. Specifically, organic calcium substrates like lactate optimize the bioconversion capacity of bacteria—but not fungi—within concrete aggregates (70, 71). In Furthermore, acidic groups within the extracellular polymeric substances (EPS) act as an organized electrostatic trap, binding free divalent cations to overcome the initial thermodynamic barrier (72, 73). However, whether this crystal initiation occurs via passive extracellular nucleation or requires active intracellular processing remains under debate (26).

By positioning our growth and consolidation data alongside these established metabolic baselines, we clarify that while the physical chemistry of acetate consumption and EPS ion-trapping follows predicted pathways, the upstream genetic circuitry—specifically the distinct bioavailability thresholds required to activate the *sinI* matrix machinery captured by our kinetic framework —represents a critical regulatory checkpoint governing the transition from basic metabolic maintenance to robust material engineering. Our findings extend this understanding by revealing a striking uncoupling between bacterial growth and biomineralization competence. Calcium (both CaCO_3_ and Ca Acetate) proved to be a growth promoter beyond its direct role in precipitation. Under our conditions, calcium can function as an essential cofactor for extracellular proteases and amylases (74, 75), thereby accelerating the hydrolysis of yeast-extract components within the biomineralization medium. Furthermore, *B. subtilis* envelope features a high density of anionic teichoic and teichuronic acids. Interfacial calcium ions serve as divalent bridges that neutralize these negative charges, structurally reinforcing the cell envelope to sustain rapid cell elongation and division while preventing autolysis(76). Consequently, the dual contribution of calcium to both biomineralization and enhanced biomass yield likely stems from this critical stabilization of wall teichoic acids.

However, calcium carbonate completely failed to induce complex colony morphology (Fig. 2c and Table S2) or *de novo* calcite precipitation in the wild-type strain, and ACC could only partially restore these phenotypes (Fig. S5). This challenges the intuitive assumption that higher biomass yields greater mineralization. Instead, our data indicate that biomineralization in *B. subtilis* is not a passive byproduct of saturation but a highly regulated physiological state. As evidenced by the independence of biofilm formation from growth parameters (Figs. 1-4), specific metabolic triggers are required, which are absent when carbonate is supplied exogenously. Importantly, our data reveals a striking physiological decoupling: while *B. subtilis* effectively “mines” insoluble calcium carbonate to support robust biomass accumulation, this source fails to trigger the *sinI* expression levels required for matrix-mediated biomineralization (Fig. 6a, Fig. 6e and S7).

By utilizing the MatTek computational framework, which proved instrumental in identifying subtle physiological states through high-resolution signal processing (Supporting File 1), we demonstrated that calcium acetate significantly enhances the expression efficiency of *sinI*, the master regulator of biofilm formation. Our analysis reveals that while the acetate moiety alone is inert, the combination of soluble calcium and acetate dramatically shifts the transcription kinetics of the matrix machinery. In contrast, the failure of calcium carbonate triggers a similar response, despite its utilization for biomass accumulation, suggests that the intracellular calcium thresholds required to induce the *sinI* pathway are higher than those needed for basic metabolic maintenance. This disparity, captured with high temporal resolution by the MatTek framework, underscores that biomineralization is not a default consequence of microbial growth in the presence of calcium, but a tightly regulated response dependent on specific cation bioavailability. These data suggest that calcium acetate may function as a specific biochemical signal, likely facilitated by its distinct solubility profile and efficient intracellular uptake. This hypothesis is consistent with the intracellular localization of the sensor kinases KinA, KinD, and KinE, which initiate the phosphorelay cascade leading to the phosphorylation of the master regulator Spo0A (77). Furthermore, these observations align with previous reports indicating that transcriptional responses within biomineralization media are critically dependent on KinA and KinB activity (53).

From an engineering perspective, the inability of wild-type *B. subtilis* to mineralize exogenous calcium carbonate (CaCO_3_; Fig. 2c and Table S2) presents a significant hurdle for biotechnologies that aim to utilize recycled construction waste as a primary feedstock. While our data suggests that the bacteria can effectively “mine” crystalline CaCO_3_ to support metabolic growth, the native regulatory circuits fail to translate this intake into the robust ECM production required for mineral nucleation.

However, our experiments with matrix-overproducing mutants demonstrate that this limitation is fundamentally regulatory and structural, rather than chemical. By genetically forcing the overproduction of exopolysaccharides (EPS) and amyloid fibers, we partially restored the ability of *B. subtilis* to form integrated mineralized structures even in the presence of poorly soluble carbonate sources (Fig. 7a). This finding presents a targeted synthetic biology strategy to bypass native environmental sensing constraints (Fig. 7b-d). This approach contributes to the development of sustainable construction technologies, moving toward actively programmed Engineered Living Materials (ELMs) rather than relying solely on passive bacterial additives.

