## Supporting Information for "Assessing soluble and insoluble calcium sources for growth, biofilm formation, and biomineralization in *Bacillus subtilis*"

**Supporting Information File**

Supporting Tables (Tables S1-2)

Supporting Figures (Figures S1-S7)

Supporting References

| Strain | Genotype / Description | References |
|---|---|---|
| *NCIB 3610* | Wild-type Bacillus subtilis , undomesticated strain capable of robust biofilm formation | **Branda et al., 2001** |
| *ΔsinR* | Derivative of NCIB 3610; deletion of the biofilm repressor *sinR* | **Kearns et al., 2005** |
| *ΔabrB* | Derivative of NCIB 3610; deletion of the transition state regulator *abrB* | **Kearns et al., 2005** |
| *amyE::*$p_{sinI}$*luciferase* | Derivative of NCIB 3610 ; *amyE*::$p_{sinI}$luciferase $Cam^R$ , transcriptional reporter for biofilm induction | **Bucher et al., 2016** |
| *ΔtasA::kan, ΔepsH*::tet | Derivative of NCIB 3610; *ΔtasA::kan, ΔepsH*::tet | **Steinberg et al., 2020** |

**Table S1. Bacterial strains used in this study.** List of *Bacillus subtilis* strains, including wild-type (NCIB 3610) and derivatives, with their respective genotypes and sources utilized for biofilm and biomineralization assays.

| Calcium source | | Day 4 | Day 10 | Day 14 | Day 18 | Day 20 | Day 24 |
|---|---|---|---|---|---|---|---|
| Calcium acetate | Dry-weight | Under DL | Under DL | Under DL | 0.00181g | 0.00207g | 0.00036g |
| | FTIR assignment of calcite | - | - | - | yes | yes | yes |
| Calcium carbonate | Dry-weight | Under DL | Under DL | Under DL | Under DL | Under DL | Under DL |

**Table S2. Temporal analysis of mineral precipitation dry weight over 24 days.** Significant accumulation of crystalline material is observed exclusively in the presence of Calcium Acetate, whereas Calcium Carbonate yields negligible precipitate below the detection limit (DL).

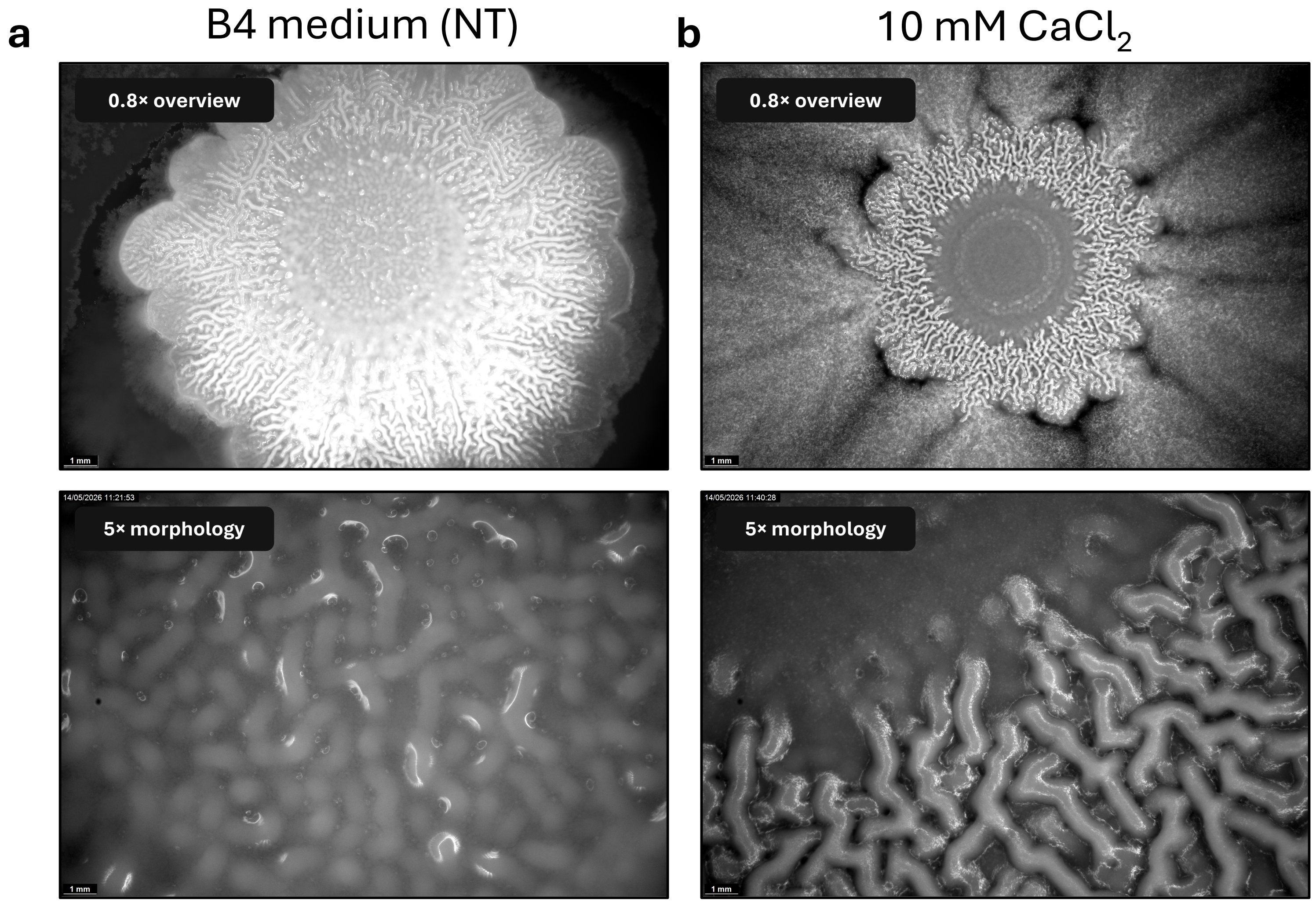


**Fig. S1. Calcium chloride affects colony morphology differently than calcium acetate.** Representative overview and magnified images of *B. subtilis* colonies grown on B4 medium without treatment (NT) or supplemented with 10 mM calcium chloride.

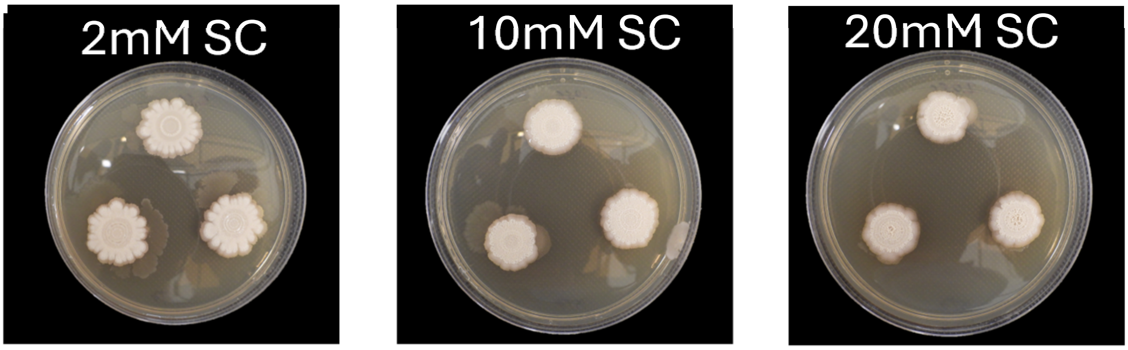


**Fig. S2. Sodium carbonate does not induce mineral-associated colony morphology.** Representative images of *B. subtilis* colonies grown on B4 medium without treatment (NT) or supplemented with sodium carbonate (SC; 2, 10, or 20 mM). Sodium carbonate did not reproduce the calcium-acetate-associated pigmented or structured colony morphology. The results represent one out of three independent experiments performed with five technical repeats. The control for this experiment is included within main figure 1.

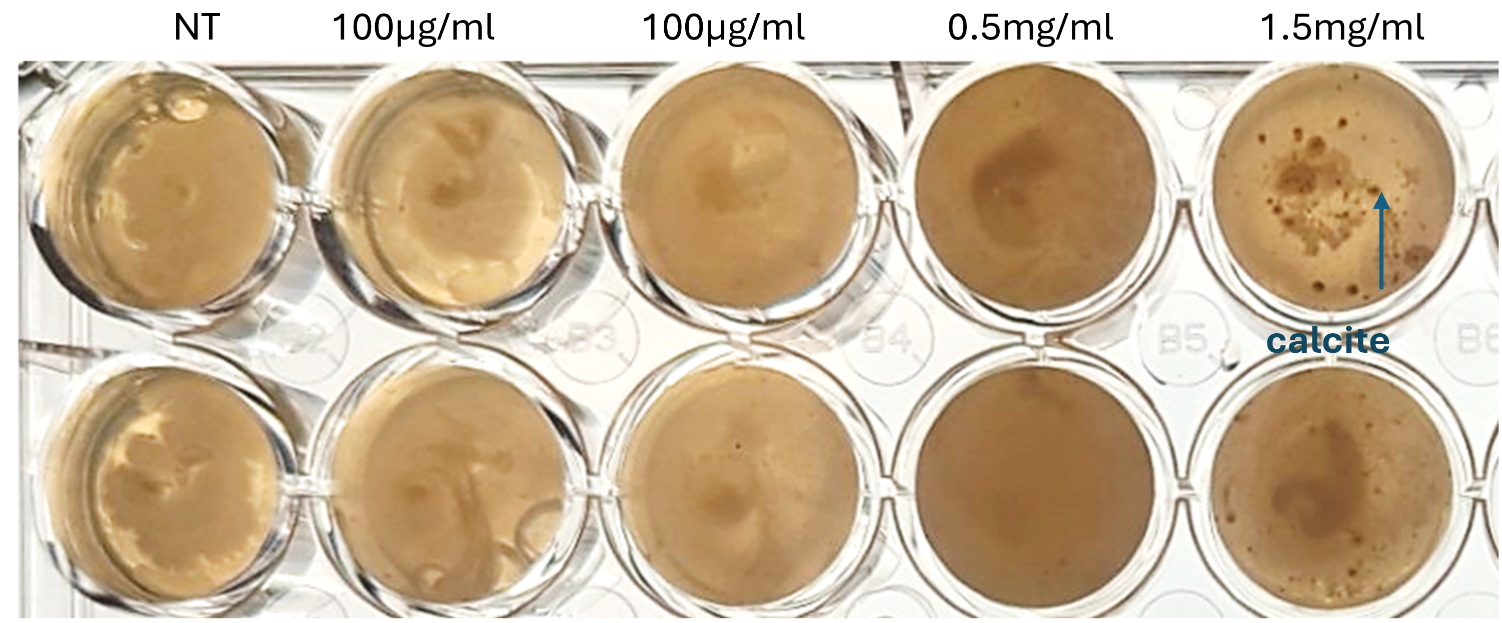


**Figure S3. Amorphous calcium carbonate induces biomineralization only at high concentrations.** Representative cultures supplemented with amorphous calcium carbonate (ACC) or a calcite reference, as indicated. Visible calcite-like precipitates were detected only at the highest ACC concentration tested, consistent with the requirement for saturated or near-saturated ACC conditions to induce mineral accumulation. Plates were images following 20 days of incubation. The results represent one out of two independent experiments performed with three technical repeats.

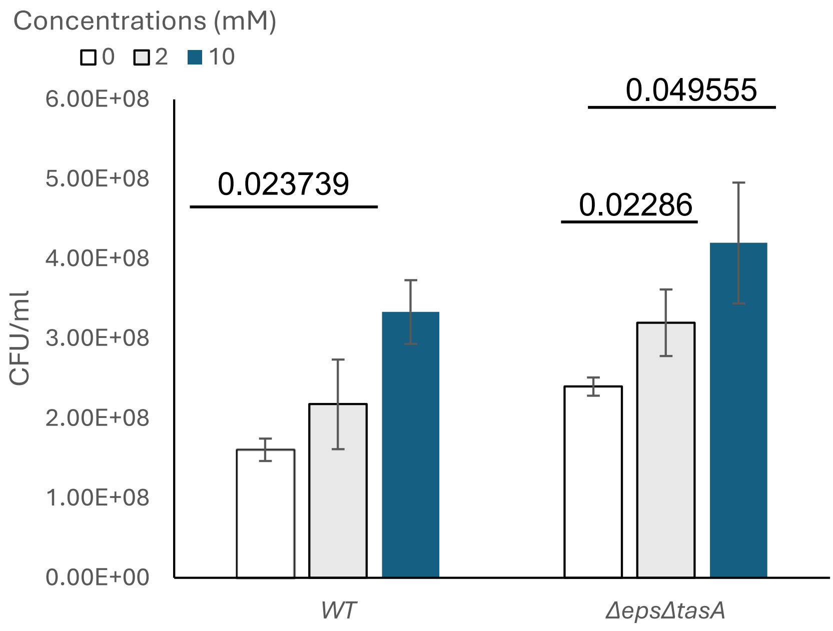


**Figure S4. Calcium acetate increases viable cell counts in *B. subtilis* planktonic cultures.** Colony-forming units (CFU/mL) recovered from wild-type *B. subtilis* and the matrix-deficient mutant *ΔepsHΔtasA* after growth on B4 medium supplemented with calcium acetate at 0, 2, or 10 mM for 8 hours. Calcium acetate increased recoverable CFU in both genetic backgrounds, with the strongest effect observed at 10 mM. Bars represent the mean ± SD. Statistical comparisons are indicated above the bars (paired t-test). The results represent the average of three independent experiments performed with two technical repeats.

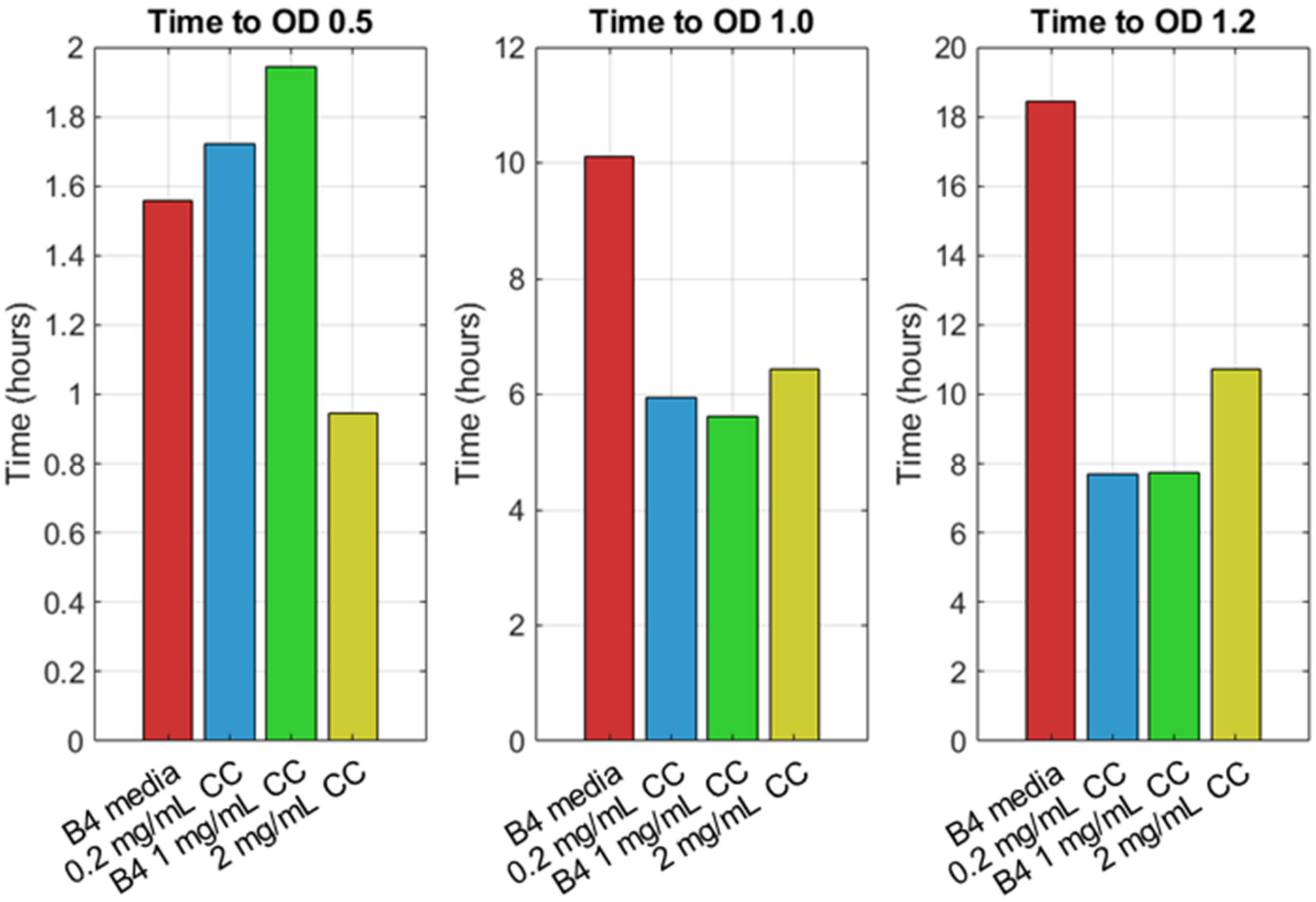


**Figure S5. Calcium carbonate supports late-stage growth in time-to-threshold analysis.** Time-to-threshold analysis showing the time required for cultures grown with insoluble calcium carbonate (CC; 0.2, 1, or 2 mg/mL) to reach $OD_{600}$ values of 0.5, 1.0, and 1.2. CC-treated cultures reached higher $OD_{600}$ thresholds faster than the B4 control at later growth stages.

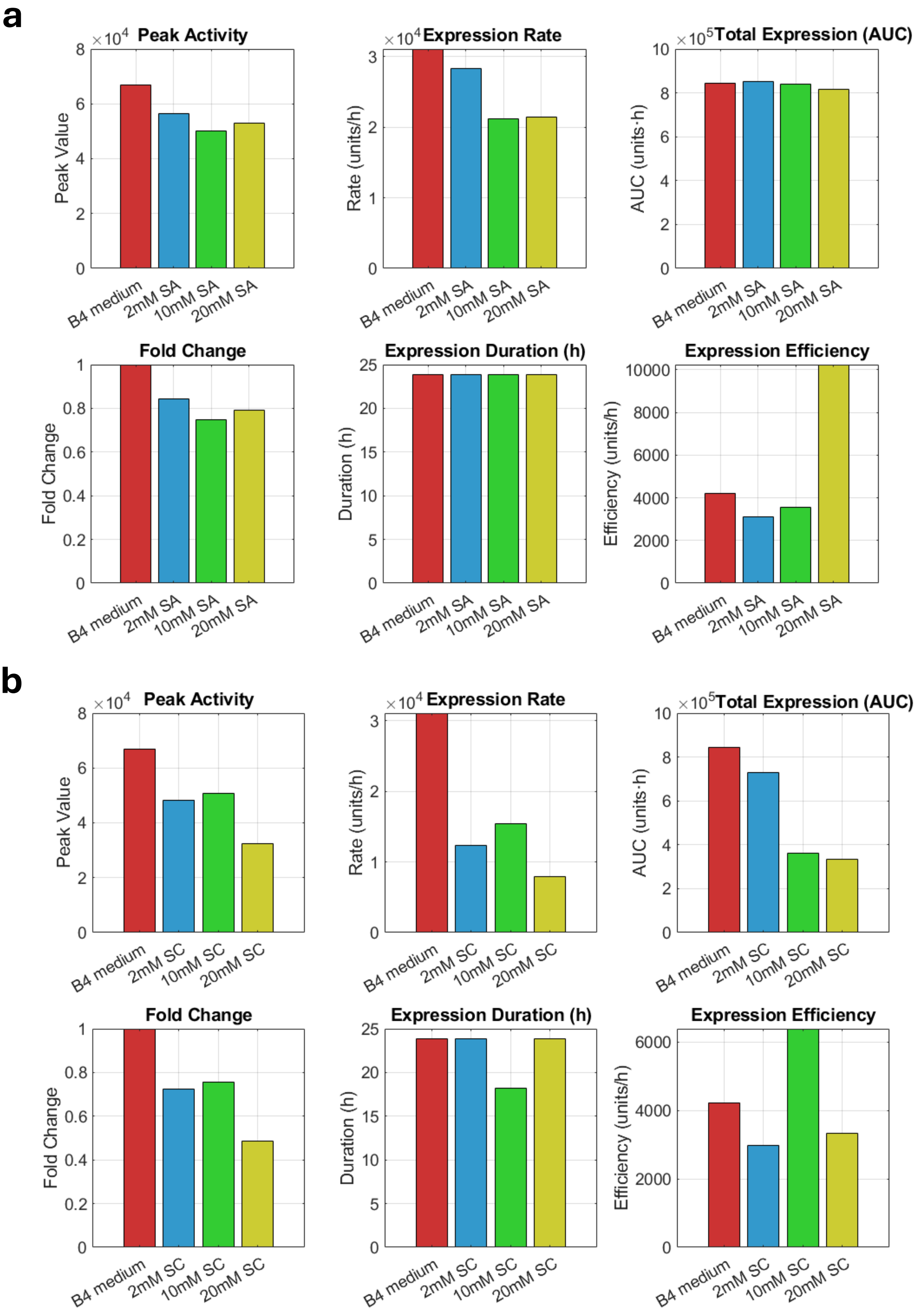


**Figure S6. Sodium salts differentially affect *sinI* reporter-expression metrics.** MatTek-derived P*sinI*-lux reporter metrics for cultures grown with (a) sodium acetate (SA; 2, 10, or 20 mM) and (b) sodium carbonate (SC; 2, 10, or 20 mM). Metrics include peak activity, expression rate, total expression (AUC), fold change, expression duration, and expression efficiency. Sodium carbonate reduced several reporter-output parameters relative to B4 medium, whereas sodium acetate produced comparatively modest changes

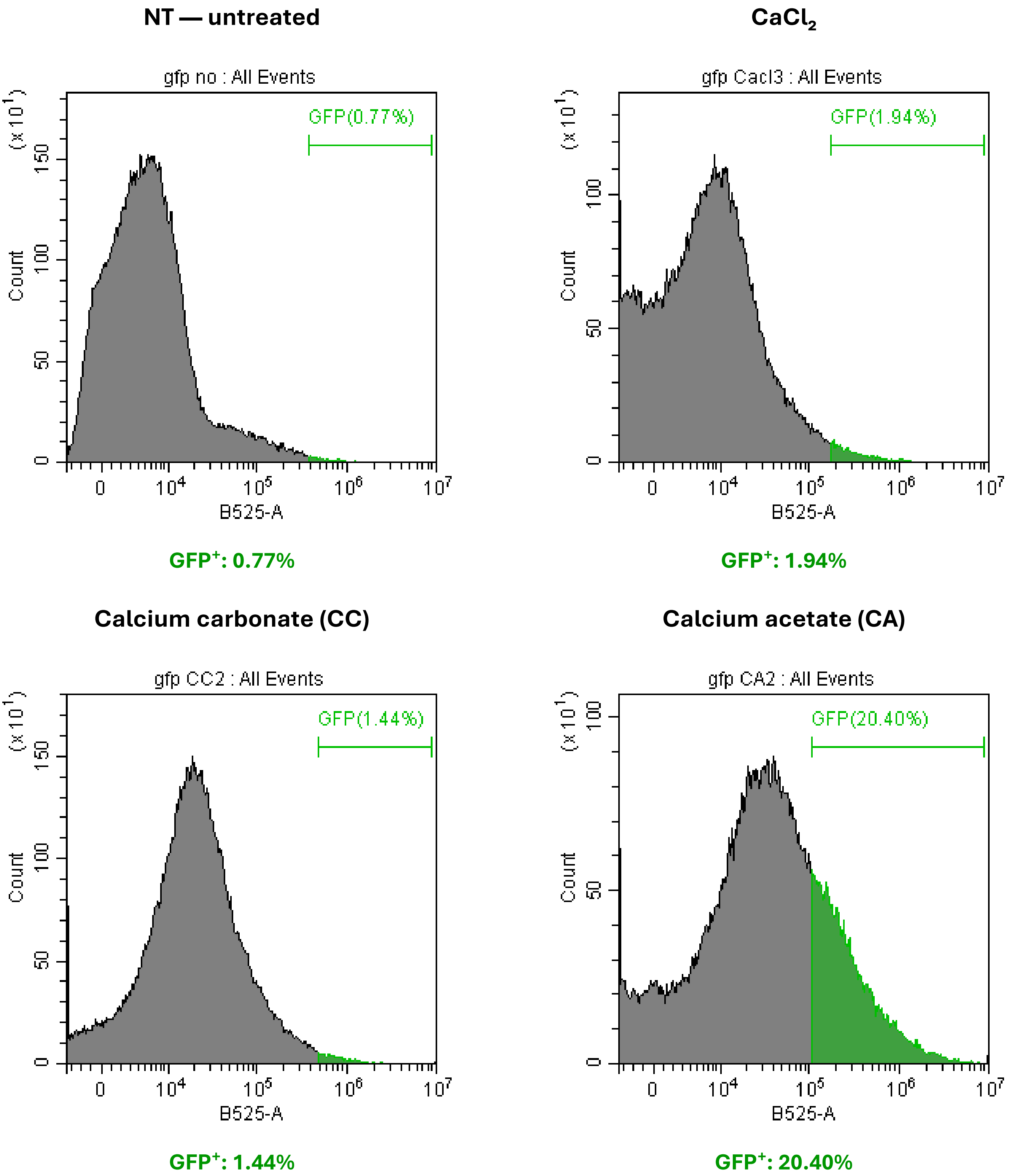


**Fig. S7. Calcium acetate selectively expands the *sinI*-GFP-positive population.** Flow-cytometry analysis of cells recovered from B4 agar plates supplemented with calcium acetate, calcium carbonate, or calcium chloride. Calcium acetate produced the largest increase in the GFP-positive population, whereas calcium carbonate and calcium chloride remained close to the untreated condition. The results represent one out of two independent experiments performed with three technical repeats.
